# The claudin-like apicomplexan microneme protein is required for gliding motility and infectivity of *Plasmodium* sporozoites

**DOI:** 10.1101/2022.08.29.505663

**Authors:** Manon Loubens, Carine Marinach, Clara-Eva Paquereau, Soumia Hamada, Bénédicte Hoareau-Coudert, David Akbar, Jean-François Franetich, Olivier Silvie

## Abstract

Invasion of host cells by apicomplexan parasites such as *Toxoplasma* and *Plasmodium* spp requires the sequential secretion of the parasite apical organelles, the micronemes and the rhoptries. The claudin-like apicomplexan microneme protein (CLAMP) is a conserved protein that plays an essential role during invasion in *Toxoplasma gondii* tachyzoites and *Plasmodium falciparum* merozoites. CLAMP is also expressed in *Plasmodium* sporozoites, the mosquito-transmitted forms of the malaria parasite, but its role in this stage is still unknown. CLAMP is essential for *Plasmodium* blood stage growth and is refractory to conventional gene deletion. To circumvent this obstacle and study the function of CLAMP in sporozoites, we used a conditional genome editing strategy based on the dimerisable Cre recombinase in the rodent malaria model parasite *P. berghei*. We successfully deleted *clamp* gene in *P. berghei* transmission stages and analyzed the functional consequences on sporozoite infectivity. In mosquitoes, sporozoite development and egress from oocysts was not affected in conditional mutants. However, invasion of the mosquito salivary glands was dramatically reduced upon deletion of *clamp* gene. In addition, CLAMP-deficient sporozoites were impaired in cell traversal and productive invasion of mammalian hepatocytes. This severe phenotype was associated with major defects in gliding motility and with reduced shedding of the sporozoite adhesin TRAP. These results demonstrate that CLAMP is essential across invasive stages of the malaria parasite, and strongly suggest that the protein acts upstream of host cell invasion, possibly by regulating the secretion or function of adhesins in *Plasmodium* sporozoites.

**Author summary:** *Plasmodium* parasites, the causative agents of malaria, are transmitted during the bite of an infected mosquito. Infectious parasite stages known as sporozoites are released from the insect salivary glands and injected into the host skin. Sporozoites rapidly migrate to the host liver, invade hepatocytes and differentiate into the next invasive forms, the merozoites, which invade and replicate inside red blood cells. Sporozoite motility and host cell invasion rely on the secretion of apical organelles called micronemes and rhoptries. Here we characterize the function of a microneme protein expressed both in merozoites and sporozoites, the claudin-like protein CLAMP. We used a conditional genome editing strategy in a rodent malaria model to generate CLAMP-deficient sporozoites. In the absence of CLAMP, sporozoites failed to invade mosquito salivary glands and mammalian hepatocytes, and showed defects in gliding motility and microneme secretion. Our data establish that CLAMP plays an essential role across *Plasmodium* invasive stages, and might represent a potential target for transmission-blocking antimalarial strategies.

## Introduction

The life cycle of *Plasmodium* parasites is complex and alternates between a vertebrate host and an *Anopheles* vector, in which parasites face several stages of differentiation and development. In mammals, infection begins when motile forms of the parasite known as sporozoites are injected in the skin during the bite of a blood-feeding infected mosquito. These sporozoites then migrate through the dermis until they reach a blood vessel and are then transported in the blood flow to the liver. There, sporozoites infect hepatocytes and differentiate into thousands of merozoites that are then released in the blood and invade red blood cells (RBCs). Sporozoites are formed in oocysts in the mosquito midgut, and once released in the haemolymph, they colonize the insect salivary glands in order to permit transmission upon salivation during bite.

Host cell invasion by *Plasmodium* and related apicomplexans relies on the sequential secretion of apical secretory vesicles, the micronemes and the rhoptries, and the formation of a structure called moving junction (MJ), which mediates parasite internalization and the formation of a replicative niche, the parasitophorous vacuole (PV). The MJ has been well characterised during host cell invasion by *Toxoplasma* tachyzoites and *Plasmodium* merozoites (Besteiro et al., 2011; Cowman et al., 2017). In contrast, the nature of the junction during sporozoite entry into hepatocytes remains elusive (Bargieri et al., 2014). Known components of the MJ, such as AMA1 and RONs, are also expressed in sporozoites, and are required during invasion not only of hepatocytes but also of the mosquito salivary glands (Fernandes et al., 2022; Ishino et al., 2019; Nozaki et al., 2020). A genome-wide CRISPR screen performed in *T. gondii* has identified a novel putative component of the MJ, the Claudin-Like Apicomplexan Microneme Protein (CLAMP) (Sidik et al., 2016). CLAMP orthologs are present in all available apicomplexan genomes and no related sequence could be identified outside the phylum. However, topology predictions indicate structural similarities between CLAMP and the mammalian tight-junction proteins claudin-15 and claudin-19 (Sidik et al., 2016). CLAMP is found in *Toxoplasma* tachyzoite micronemes and at the MJ during invasion of host cells. Conditional knockdown of *clamp* gene affects *T. gondii* host cell invasion without disturbing egress or microneme secretion (Sidik et al., 2016). Similarly, depletion of CLAMP in *P. falciparum* merozoites leads to a drastic reduction in RBC invasion (Sidik et al., 2016). CLAMP is also predicted to be essential in *P. berghei* based on genome-wide mutagenesis (Bushell et al., 2017). A more recent study focusing on the role of CLAMP in *Theileria equi* showed that CLAMP-specific antibodies inhibit invasion of equine erythrocytes (Onzere et al., 2021).

While proteomic studies have documented that CLAMP is also expressed in sporozoites in multiple *Plasmodium* species (Hamada et al., 2021; Lindner et al., 2013; Swearingen et al., 2017), its role has never been assessed so far in these stages. One obstacle for genetic studies of factors like CLAMP is that genes that are essential in blood stages, the stages that are used for genetic manipulation, are refractory to conventional knock-out strategies. Nevertheless, several approaches for conditional gene deletion have emerged allowing the study of blood stage-essential genes in sporozoites. One of these approaches is based on the dimerisable Cre recombinase (DiCre), which can excise DNA sequences flanked by Lox sites in an inducible manner. The recombinase is split into two inactive subunits each fused to a rapamycin-ligand, which interact in the presence of rapamycin, restoring the Cre activity (Andenmatten et al., 2012; Jullien et al., 2003). *Plasmodium* asexual blood stage parasites can be targeted prior transmission to mosquitoes, allowing deletion of the gene of interest and phenotypical analysis in subsequent stages of the parasite life cycle (Fernandes et al., 2020; Tibúrcio et al., 2019). In this study, we used the DiCre system in *P. berghei* to study the role of CLAMP in sporozoites. Conditional deletion of *clamp* gene resulted in a dramatic decrease of sporozoite invasion of the mosquito salivary glands and of mammalian hepatocytes. This severe phenotype was associated with a major defect in sporozoite gliding motility and reduced shedding of TRAP. This study reveals that CLAMP is required for *Plasmodium* sporozoite motility and infectivity in both the mosquito and mammalian hosts, and illustrates the robustness of the DiCre system for conditional genome editing across the parasite life cycle.

## Results

### Conditional deletion of *clamp* gene in *P. berghei*

In order to study the role of CLAMP in *P. berghei* sporozoites, we first engineered a parasite line allowing conditional disruption of *clamp* gene using the DiCre system. We genetically modified a *P. berghei* line (PbDiCre) expressing the two DiCre components in addition to a mCherry fluorescence cassette (Fernandes et al., 2020) in order to generate a *clamp*cKO line where the CLAMP-coding sequence is fused to a 3xFlag tag and flanked by LoxN sites. For this purpose, we used a two-step strategy to introduce in two successive transfections LoxN sites upstream and downstream of *clamp* gene, respectively **(Figs 1A** and **S1)** (Fernandes et al., 2021). A GFP cassette was introduced between *clamp* and the second LoxN site, to facilitate monitoring of gene excision by flow cytometry or fluorescence microscopy. Transfected parasites were selected with pyrimethamine and sorted by flow cytometry, and the final *clamp*cKO parasites were cloned by limiting dilution and injection into mice. The parasites were genotyped by PCR to confirm correct construct integration **(S1 Fig)**. Transfected parasites were monitored by flow cytometry during the different selection steps in order to confirm the presence or absence of the GFP cassette (**Fig 1B**).

**Fig 1.**
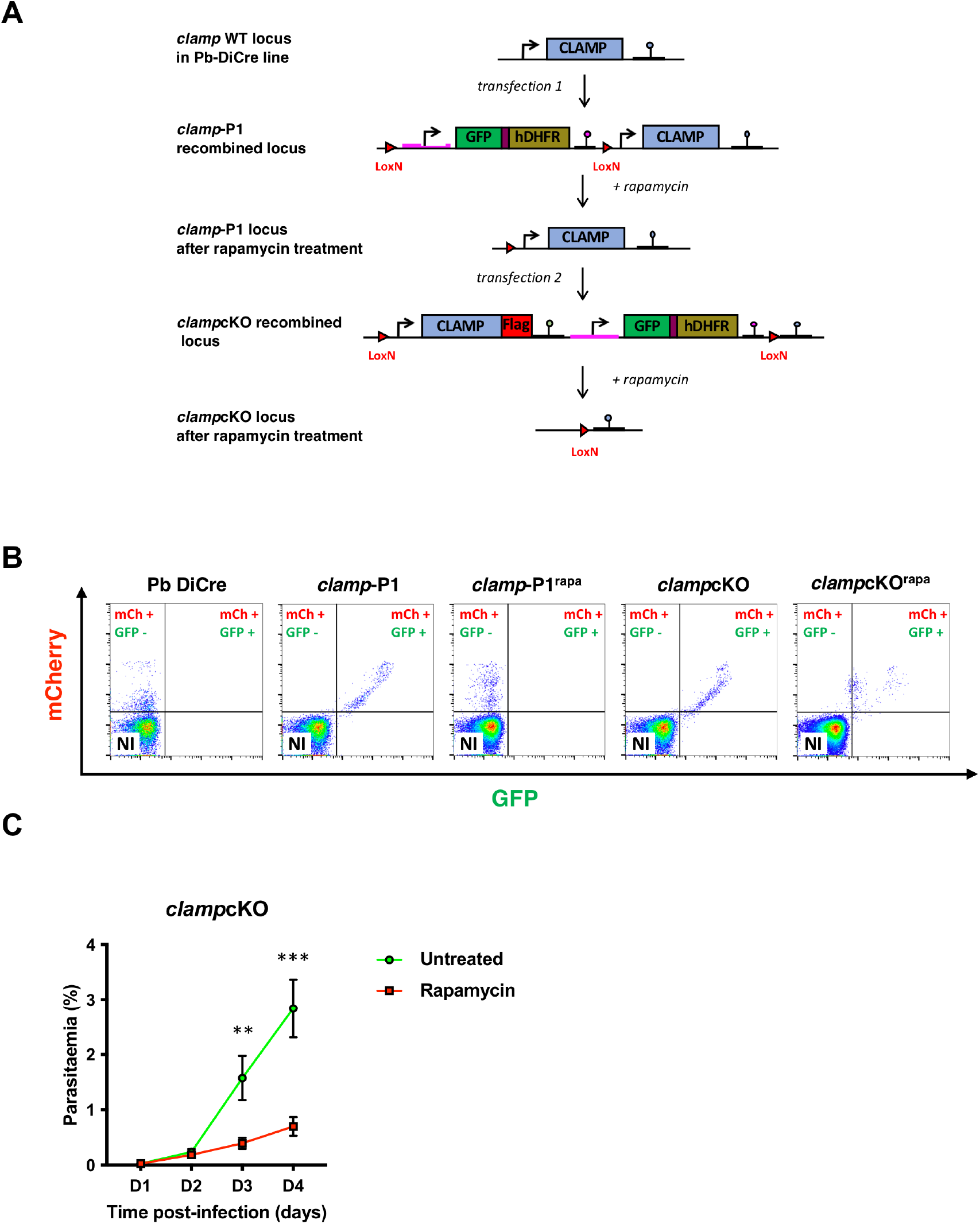
Generation of *P. berghei clamp*cKO parasites. **A**. Strategy to modify the *clamp* locus in the PbDiCre line. A first transfection in PbDiCre parasites with the P1 construct led to the insertion of a GFP-hDHFR expression cassette flanked by two LoxN site upstream of *clamp* gene. Rapamycin-induced excision of the cassette caused retention of one single LoxN site upstream of *clamp* in the treated *clamp*-P1 parasites. In a second step, rapamycin-treated excised *clamp*-P1 parasites were transfected with the P2 construct to generate a *clamp*cKO parasite line expressing CLAMP fused to a 3xFlag epitope tag in addition to GFP and hDHFR, and carrying two LoxN sites flanking *clamp* gene. Rapamycin-induced activation of DiCre leads to excision of the *clamp* gene together with the GFP-hDHFR cassette. **B**. Flow cytometry analysis of blood-stage parasites at different steps after initial transfection of the PbDiCre parental line to generate *clamp*cKO parasites. NI, non-infected red blood cells. **C**. Blood stage growth of rapamycin-treated and untreated *clamp*cKO parasites. Rapamycin was administered at day 1. The graph shows the parasitaemia (mean +/-SEM) in groups of 5 mice, as quantified by flow cytometry based on mCherry detection. **, p < 0.01; ****, p < 0.0001 (Two-way ANOVA).

We then assessed the effects of rapamycin on *clamp*cKO parasites during blood-stage growth. Following intravenous injection of 10^6^ parasitized RBCs into mice, one cohort was treated with a single oral dose of rapamycin, while the other was left untreated. We then monitored the parasitaemia over time by flow cytometry. Rapamycin exposure led to a decrease of GFP fluorescence in *clamp*cKO parasites, confirming the efficiency of gene excision (**Fig 1B**), yet we fail to recover mCherry^+^/GFP^-^ parasite populations. Rapamycin-induced excision of *clamp* greatly reduced blood-stage growth in *clamp*cKO-infected mice, as compared to untreated parasites (**Fig 1C)**, in agreement with an essential role for CLAMP in asexual blood stages (Bushell et al., 2017; Sidik et al., 2016).

### CLAMP is required for invasion of mosquito salivary glands

We next investigated the role of CLAMP in *P. berghei* mosquito stages. For this purpose, mosquitoes were fed on *clamp*cKO-infected mice, which were treated or not with rapamycin one day prior blood feeding, as described (Fernandes et al., 2020). We then monitored parasite development in the mosquito midgut and colonization of the insect salivary glands using fluorescence microscopy. Analysis of dissected midguts at day 16 post-feeding showed that both untreated and rapamycin-exposed parasites produced mCherry-positive oocysts **(Fig 2A)**. As expected, only untreated parasites retained the GFP fluorescence, while oocysts from rapamycin-treated parasites were GFP-negative, confirming efficient gene excision **(Fig 2A)**. Mosquito salivary glands dissected at day 21 post-feeding exhibited a strong fluorescence with both mCherry and GFP, reflecting sporozoite accumulation in the glands **(Fig 2B)**. Strikingly, salivary glands from rapamycin-exposed *clamp*cKO-infected mosquitoes showed a weak mCherry fluorescence signal, suggesting low parasite loads **(Fig 2B)**. Sporozoites were then isolated from midguts or salivary glands and counted. Rapamycin treatment of *clamp*cKO parasites had no major impact on the number of midgut sporozoites **(Fig 2C)**, but severely reduced the number of salivary gland sporozoites **(Fig 2D)**. As expected, rapamycin treatment induced robust gene excision in *clamp*cKO parasites, as determined based on the percentage of excised (mCherry^+^/GFP^-^) and non-excised (mCherry^+^/GFP^+^) sporozoites **(S2A** and **2S2B Figs)**. Reduced numbers of salivary gland sporozoites could result either from a defect in parasite egress from oocyst or from a defect in salivary gland invasion. We isolated and counted haemolymph sporozoites in rapamycin-exposed and untreated *clamp*cKO-infected mosquitoes and observed no significant difference **(Fig 2E)**. Mosquitoes infected with parasites expressing mCherry exhibit a red fluorescence of pericardial cells following uptake of sporozoites released in the haemolymph (Fernandes et al., 2022). A large majority of infected mosquitoes presented such mCherry-labelled structures, with both rapamycin-exposed and untreated *clamp*cKO parasites **(S2C Fig)**, indicating efficient egress from oocysts in CLAMP-deficient parasites. Together, these results show that sporozoites deleted for *clamp* are able to develop normally in the mosquito until release in the haemolymph but are then impaired in their ability to invade the salivary glands.

**Fig 2.**
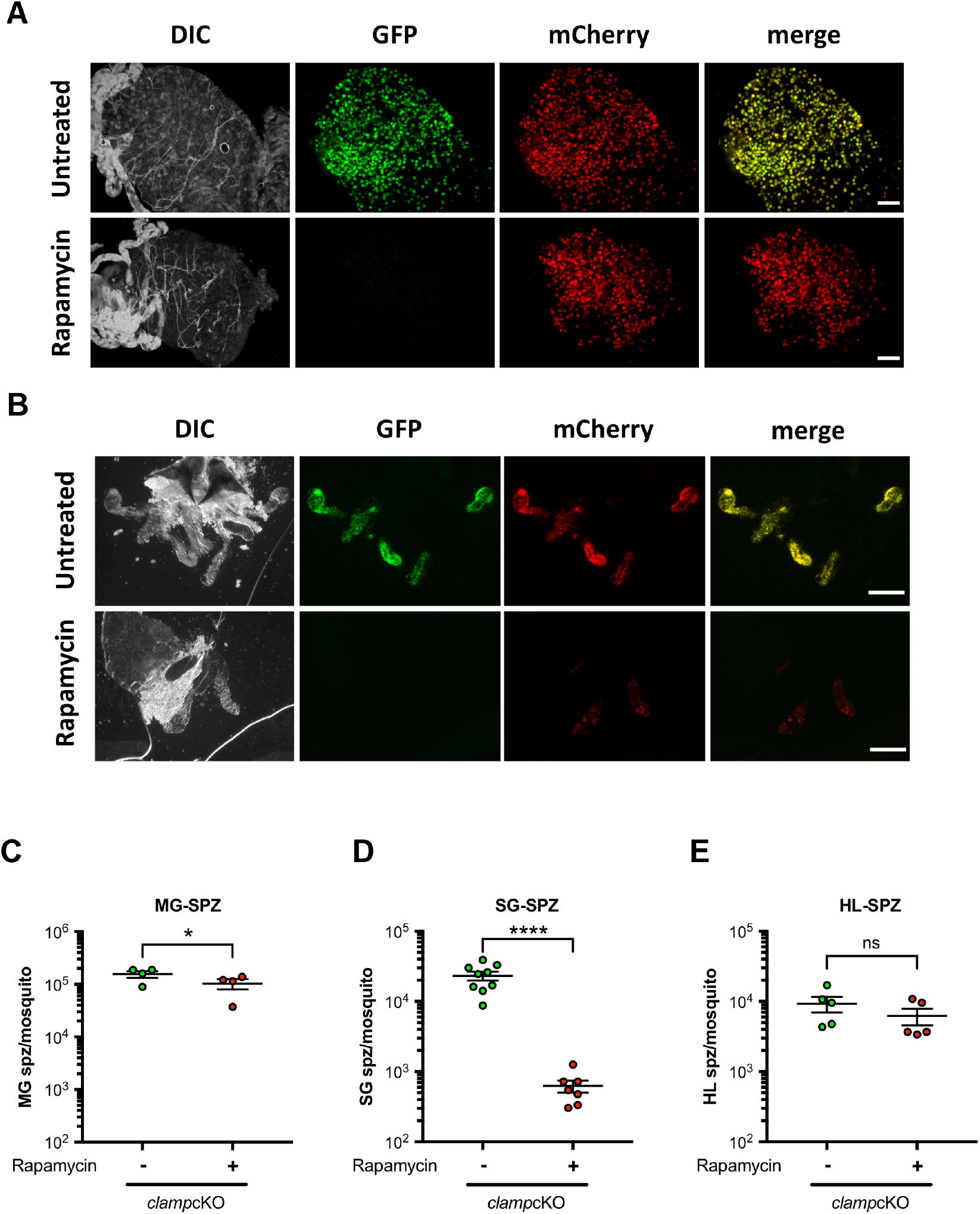
Deletion of *clamp* impacts invasion of mosquito salivary glands by sporozoites. **A-B**. Fluorescence microscopy imaging of midguts (A) and salivary glands (B) from rapamycin-exposed and untreated *clamp*cKO-infected mosquitoes, dissected at day 16 or 21 post-feeding, respectively. Exposure time and contrast were adjusted at the same level for each channel in both conditions. Scale bar, 200 μm. **C-E**. Comparison of sporozoite numbers isolated from midguts (C), salivary glands (D) and haemolymph (E) of female mosquitoes infected with rapamycin-exposed and untreated *clamp*cKO parasites. All results shown in C-E are mean +/-SEM of at least four independent experiments. Ns, non-significant; *, p < 0.05; ****, p<0.0001 (Two-tailed ratio paired t test).

### Rapamycin-induced gene excision abrogates CLAMP protein expression in sporozoites

In order to verify that *clamp* gene excision induced by rapamycin exposure was efficient in depleting CLAMP protein, we took advantage of the 3xFlag fused to the C-terminus of CLAMP in the *clamp*cKO line (**Fig1A**). Analysis of untreated *clamp*cKO salivary gland sporozoites by immunofluorescence using anti-Flag antibodies revealed a granular distribution of the protein in the cytoplasm of the parasite, consistent with a localization in the micronemes **(Fig 3A)**. In contrast, no labelling was observed in rapamycin-exposed *clamp*cKO sporozoites, confirming complete depletion of the protein upon *clamp* gene excision **(Fig 3A)**. We also confirmed by western blotting the absence of CLAMP protein in haemolymph sporozoites after rapamycin treatment, while in untreated parasites the protein was detected as a single ∼ 40 kDa band (**Fig 3B**). These data thus confirm that CLAMP is expressed in *P. berghei* sporozoites and validate the conditional approach to deplete CLAMP protein in sporozoites.

**Fig 3.**
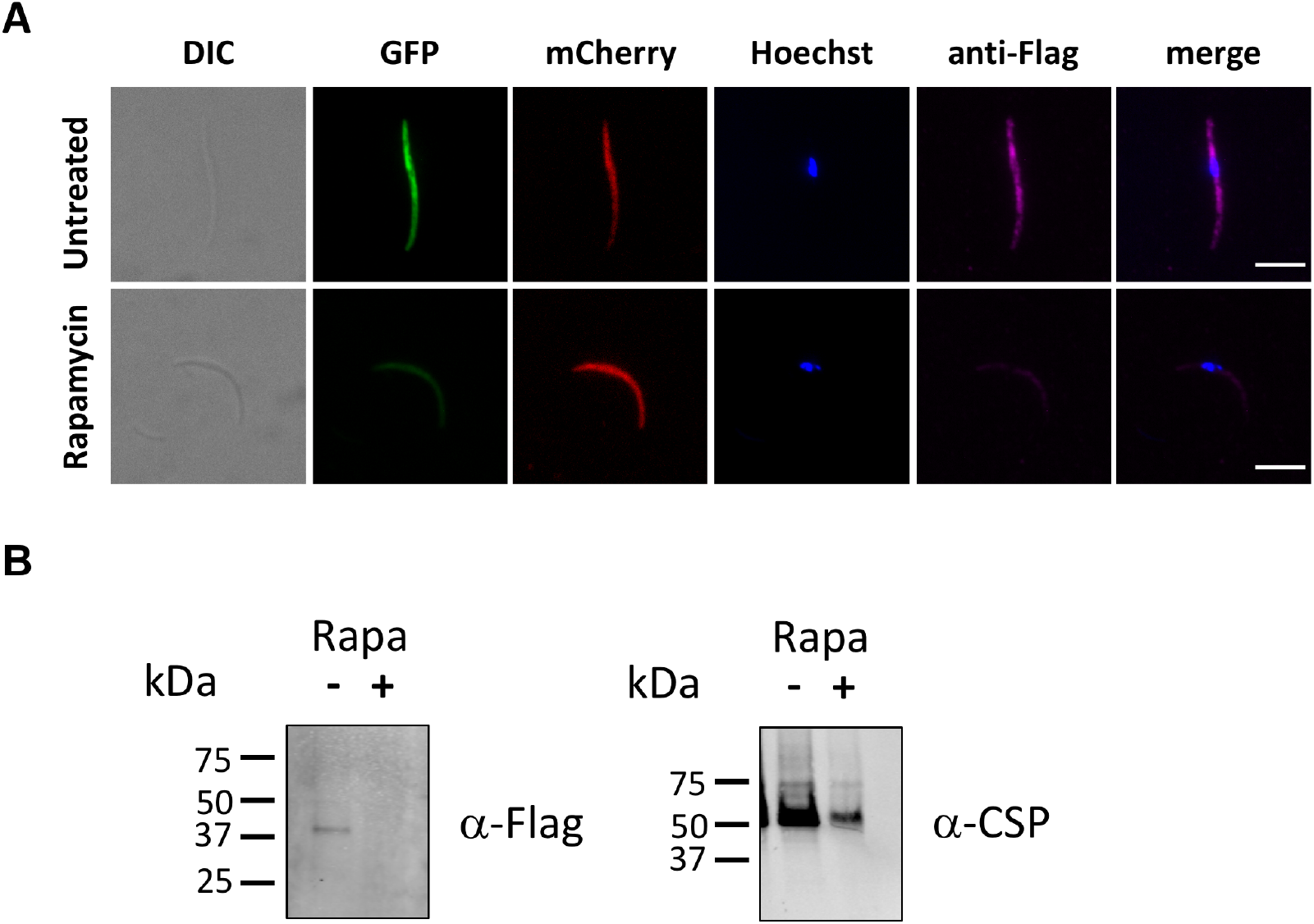
Conditional gene deletion abrogates CLAMP protein expression in sporozoites. **A**. Immunofluorescence analysis of rapamycin-exposed and untreated *clamp*cKO sporozoites labelled with anti-Flag antibodies (magenta). Both rapamycin-exposed and untreated *clamp*cKO parasites express mCherry (red), while only untreated *clamp*cKO parasites express GFP (green). Nuclei were stained with Hoechst 77742 (blue). Scale bar, 5 μm. **B**. Western bot analysis of haemolymph sporozoite lysates from untreated or rapamycin-treated parasites, using anti-Flag antibodies to detect CLAMP. CSP was used as a loading control.

### CLAMP is essential for sporozoite cell traversal and hepatocyte invasion

We then assessed the capacity of salivary gland sporozoites to traverse and invade hepatocytes *in vitro* in the absence of CLAMP. We first quantified by flow cytometry the number of traversed cells 3 hours after sporozoite addition to HepG2 cell cultures, using a dextran-based assay as previously described (Mota et al., 2001; Prudêncio et al., 2008). Cell traversal was severely impaired in sporozoites lacking CLAMP, as shown by a dramatic reduction of the percentage of dextran-positive cells in cultures incubated with rapamycin-exposed *clamp*cKO as compared to untreated *clamp*cKO sporozoites **(Fig 4A)**. We next tested if sporozoites lacking CLAMP could infect and develop into EEFs in cell cultures. HepG2 cells were incubated with rapamycin-exposed and untreated *clamp*cKO sporozoites and EEFs were quantified at 24h post-infection by microscopy after staining of PVs with antibodies against UIS4, a marker of the PV membrane (Mueller et al., 2005). These experiments revealed an almost complete absence of EEFs in cultures inoculated with rapamycin-exposed *clamp*cKO parasites **(Fig 4B)**. These data show that, in addition to its role during salivary gland invasion in the mosquito, CLAMP is also essential in sporozoites for cell traversal and productive invasion of mammalian cells.

**Fig 4.**
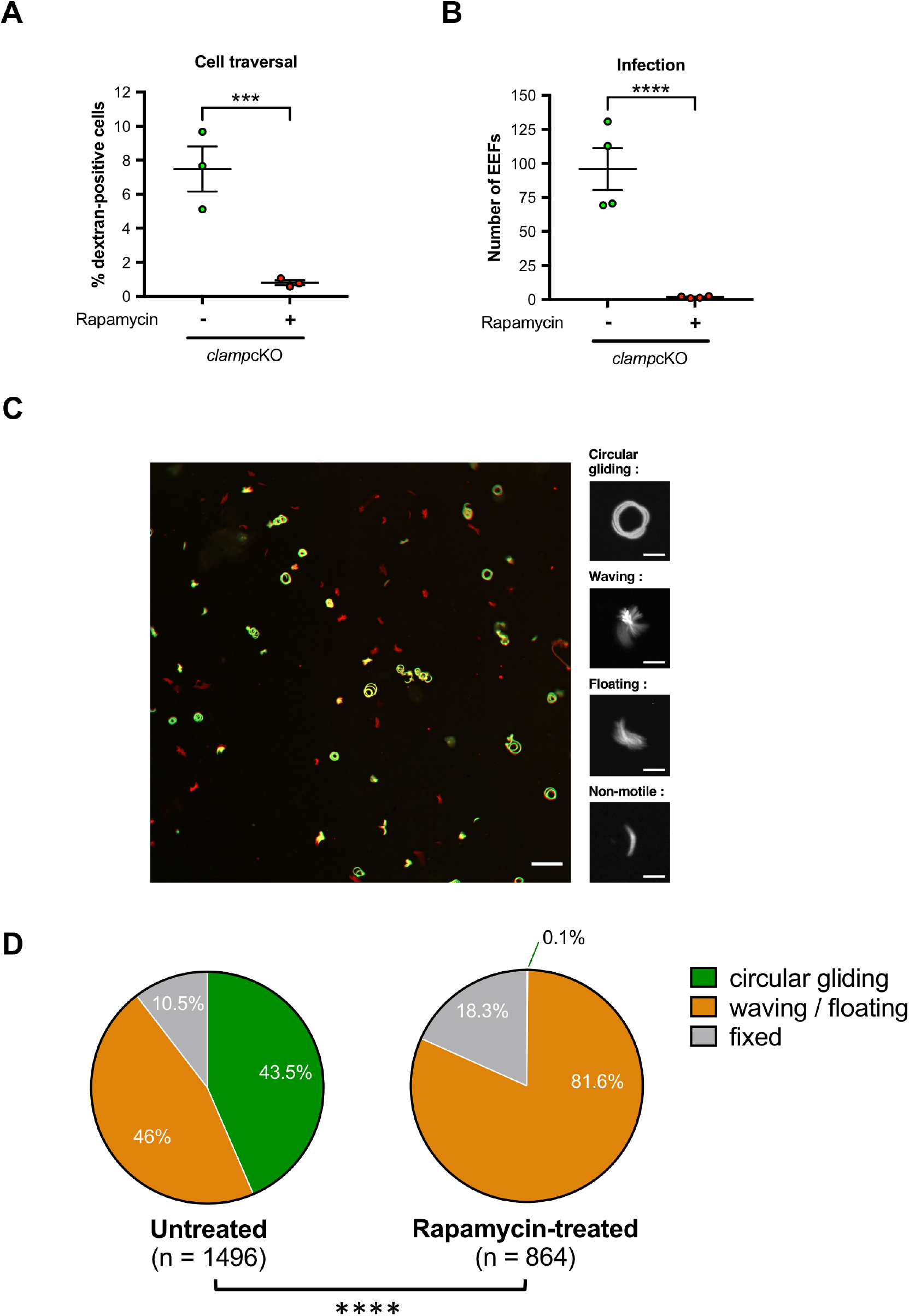
CLAMP-deficient sporozoites are impaired in transcellular migration, host cell infection and motility. **A**. Quantification of traversed (dextran-positive) HepG2 cells by FACS after incubation for 3h with rapamycin-exposed and untreated *clamp*cKO sporozoites in the presence of fluorescein-labelled dextran. Results shown are mean +/-SEM of three independent experiments, each performed with 5 technical replicates. **B**. Quantification of UIS4-labelled exo-erythrocytic forms (EEFs) in HepG2 cells as determined by fluorescence microscopy 24h post-invasion with rapamycin-exposed and untreated *clamp*cKO sporozoites. Results shown are mean +/-SEM of four independent experiments with 5 technical replicates for each. ***, p<0.001; ****, p<0.0001 (Two-tailed ratio paired t test). **C**. Maximum intensity projection of video-microscopy images of untreated *clamp*cKO (mCherry^+^/GFP^+^, yellow) and rapamycin-exposed *clamp*cKO (mCherry^+^/GFP^-^, red) sporozoites, recorded for 3 min. Sporozoites were mixed and activated at 37°C in the presence of albumin before imaging. Their movement patterns were classified in four categories : circular gliding, waving, floating and fixed (shown in magnified images). Scale bars, 50 μm for projection and 10 μm for magnified images. **D**. Quantification of motility patterns in untreated (n = 1496) and rapamycin-exposed (n = 864) *clamp*cKO sporozoites. Sporozoites were classified in three groups based on their motility pattern. ****, p<0.0001 (Chi-square).

### CLAMP is essential for sporozoite gliding motility

Since cell traversal and productive host cell invasion both rely on the parasite gliding machinery, we hypothesized that the combined phenotype observed with parasites lacking CLAMP could be caused by a defect in sporozoite motility. To address this hypothesis, we analyzed the motility of rapamycin-exposed and untreated *clamp*cKO by video-microscopy. One limitation of the assessment of sporozoite motility was the medium of dissection that was enriched in mosquito debris in the case of rapamycin-exposed parasites as compared to untreated parasites, due to the highly reduced number of salivary gland sporozoites. To ensure that the differences in the composition of the medium would not affect the motility of the parasite, we mixed untreated *clamp*cKO (mCherry^+^/GFP^+^) and rapamycin-exposed *clamp*cKO (mCherry^+^/GFP^-^) sporozoites and imaged the mixed population (**Movie 1**). Sporozoite motility was classified in three groups based on the movement patterns: circular gliding, waving or floating, or fixed (non-motile) **(Fig 4C)**. The effect of *clamp* gene deletion was striking, as over 864 observed *clamp*cKO sporozoites in the rapamycin-treated condition, only 1 (0,1%) was motile and showed a circular gliding pattern. The vast majority of rapamycin-exposed *clamp*cKO were observed waving or floating (81,6%) **(Fig 4D** and **Movie 1)**. In comparison, more than 40% of untreated *clamp*cKO sporozoites were motile and exhibited circular gliding (43,5%) **(Fig 4D** and **Movie 1)**. These results show that circular gliding is abrogated in sporozoites in the absence of CLAMP. This phenotype could explain the defects observed in salivary gland invasion and in cell traversal and hepatocyte invasion, as motility is needed by the parasite to actively enter or exit cells.

### CLAMP is required for TRAP shedding

In order to get more insights into the function of CLAMP, we searched for potential interacting partners of the protein in sporozoites through immunoprecipitation (IP) followed by mass spectrometry (MS). IP was performed using anti-Flag antibodies on two independent lysates from untreated *clamp*cKO salivary gland sporozoites. Control IP was performed using lysates from untagged PbGFP sporozoites. Analysis of samples by MS revealed 7 proteins that were identified in the two co-IP experiments but not in control IP experiments: CLAMP, the thrombospondin-related anonymous protein (TRAP, PBANKA_1349800), a putative pantothenate transporter (PAT, PBANKA_0303900), the elongation factor 1-alpha (PBANKA_1133300), the tubulin beta chain (PBANKA_1206900), actin I (PBANKA_1459300), and a conserved protein of unknown function (PBANKA_0403700) (**Fig 5A** and **S1 Table**). TRAP is a sporozoite adhesin secreted from micronemes that connects extracellular surfaces to the parasite intracellular actin-myosin motor machinery and is essential for sporozoite gliding motility (Ejigiri et al., 2012; Sultan et al., 1997). Interestingly, PAT has been shown to regulate exocytosis of osmiophilic bodies and micronemes in *P. berghei* gametocytes and sporozoites, respectively (Kehrer et al., 2016b). Sporozoites lacking PAT fail to secrete TRAP, and, as a result, are immotile and thus unable to infect host cells (Kehrer et al., 2016b). Since the phenotype of CLAMP-deficient sporozoites is reminiscent of that of TRAP- and PAT-deficient parasites, and based on our mass spectrometry data suggesting a potential interaction between these proteins, we investigated whether CLAMP could be involved in TRAP secretion. For this purpose we used haemolymph sporozoites as the number of CLAMP-deficient salivary gland sporozoites was too low to allow western blot experiments. Rapamycin-exposed and untreated *clamp*cKO haemolymph sporozoites were incubated at 37°C in the presence of albumin and ethanol to stimulate microneme secretion (Brown et al., 2020), and shedding of TRAP in the supernatant (SN) was assessed by western-blot. TRAP was detected in sporozoite lysates (pellet) from both rapamycin-exposed and untreated *clamp*cKO parasites as a main band around 100kDa, which corresponds to the expected size of the mature protein in *P. berghei* (Ejigiri et al., 2012; Klug et al., 2020). Two additional minor bands were also detected, likely reflecting different forms of the protein in haemolymph sporozoites (**Fig 5B**). A shed form of TRAP was detected in the supernatant of untreated *clamp*cKO sporozoites as a band around 75kDa, but was nearly absent in the supernatant of rapamycin-exposed *clamp*cKO sporozoites, revealing a defect of TRAP shedding in parasites lacking CLAMP (**Fig 5B**). Quantification of band intensity showed a ∼ 90% reduction of TRAP release in the supernatant as compared to sporozoite-associated protein (**Fig 5C**). These results strongly suggest that CLAMP drives sporozoite motility by regulating the secretion and/or cleavage of TRAP.

**Fig 5.**
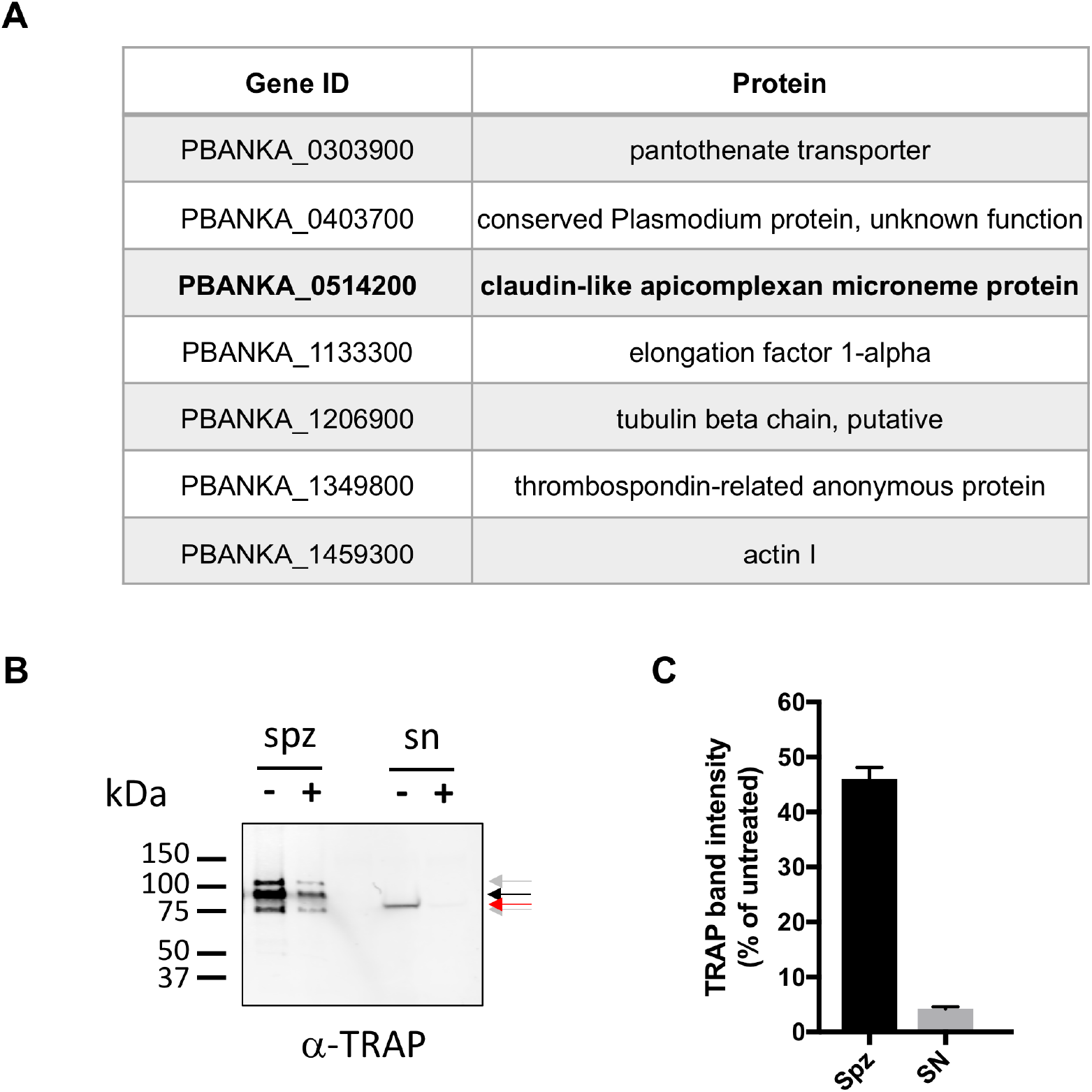
CLAMP is essential for TRAP shedding. **A**. *P. berghei* proteins identified by mass spectrometry after anti-Flag immunoprecipitation from two independent CLAMP-flag sporozoite lysates, and not from control samples. **B**. TRAP secretion assay using haemolymph sporozoites (5 × 10^4^ or equivalent per lane) from untreated or rapamycin-treated *clamp*cKO parasites. Microneme secretion was stimulated by incubation for 15 min at 37°C in the presence of BSA and ethanol. After stimulation of microneme secretion, samples were fractionated by centrifugation in sporozoite pellets (spz) and supernatants containing secreted proteins (SN), and analyzed by western blot using anti-TRAP antibodies. TRAP protein was detected as a ∼ 100 kDa major band (black arrow) and two minor bands (grey arrows) in sporozoite lysates, and as a single ∼ 75 kDa in supernatants (red arrow). **C**. Quantification of signal intensity of TRAP western blot bands in sporozoite pellets (main band only) and supernatants. The data show TRAP levels in rapamycin-treated parasites relative to untreated parasites, both in pellets and supernatants.

## Discussion

We report here the first functional characterization of CLAMP in *Plasmodium* sporozoites, revealing its crucial role in gliding motility and infectivity in both the mosquito and mammalian hosts. We generated a PbDiCre line in which *clamp* gene and a GFP expression cassette were flanked by LoxN sites. We first validated the efficient excision of *clamp* gene in blood stages, based on the disappearance of the GFP fluorescence upon exposure to rapamycin, yet we failed to establish stable CLAMP-deficient (GFP-negative) parasite populations, consistent with CLAMP being essential in blood stages (Bushell et al., 2017; Sidik et al., 2016). Transmission of CLAMP-deficient parasites to mosquitoes revealed that in the absence of the protein sporozoites develop normally and egress from oocysts, but fail to invade the salivary glands. Nevertheless, despite the low numbers of salivary gland sporozoites, we could analyze their phenotype in functional assays, which revealed that, in addition to the defect in salivary gland invasion in the mosquito, CLAMP-deficient sporozoites are impaired in both traversal and invasion of mammalian hepatocytic cells, associated with disruption of gliding motility.

The severe defect in gliding motility is likely responsible for the observed phenotype of CLAMP-deficient sporozoites, since motility is required for both cellular transmigration and host cell invasion. Sporozoite proteins specifically involved in cell traversal, such as SPECT and PLP1, differ from CLAMP as they are not required for motility, invasion of the mosquito salivary glands or productive invasion of liver cells, at least *in vitro* (Ishino et al., 2005a, 2004; Risco-Castillo et al., 2015). Reciprocally, proteins acting specifically during host cell invasion, such as the 6-cysteine domain proteins P36 and P52, are not required for gliding motility and cell traversal (Arredondo et al., 2018; Ishino et al., 2005b; Manzoni et al., 2017). CLAMP-deficient sporozoites also have a distinct phenotype as compared to parasites lacking AMA1, a canonical component of the MJ. Although in both cases sporozoites show a defect in invasion of mosquito salivary glands and mammalian hepatocytes, AMA1 conditional mutant sporozoites have no defect in gliding motility and cell traversal (Fernandes et al., 2022). Altogether, these observations strongly support a role of CLAMP upstream of host cell invasion and MJ formation in *Plasmodium* sporozoites, unlike previously proposed in *Toxoplasma* tachyzoites (Sidik et al., 2016).

Like in other apicomplexan invasive stages, sporozoite motility relies on a multimolecular machinery called the glideosome, which is localized underneath the parasite pellicle and links the parasite actin-myosin motor to surface adhesins that interact with extracellular substrates (Frénal et al., 2017). The micronemal protein TRAP is an adhesin that plays a pivotal role in sporozoite gliding motility. Disruption of *trap* gene abrogates sporozoite motility and prevents invasion of the mosquito salivary glands as well as mammalian hepatocytes (Kappe et al., 1999; Matuschewski et al., 2002; Sultan et al., 1997). We show here that CLAMP-deficient sporozoites display a similar phenotype as TRAP knockout parasites. Interestingly, TRAP was identified by mass spectrometry in CLAMP co-IP experiments, as well as the pantothenate transporter PAT, which has been shown to be required for TRAP secretion and sporozoite motility in *P. berghei* (Kehrer et al., 2016b). PAT plays an additional role in gametocytes, where it regulates exocytosis of osmiophilic bodies during egress (Kehrer et al., 2016b). Therefore, PAT is essential for parasite transmission to the mosquito (Hart et al., 2014; Kehrer et al., 2016b; Kenthirapalan et al., 2016). In our experimental setup, DiCre-mediated excision of *clamp* gene can be induced only shortly before transmission, preventing the analysis of CLAMP role in gametocytes or ookinetes.

The identification of both TRAP and PAT in co-IP experiments from salivary gland sporozoites, combined with the observed defect in TRAP shedding in CLAMP-deficient haemolymph sporozoites, strongly supports the existence of a functional link between CLAMP and TRAP-mediated gliding motility. Whether CLAMP participates in a multimolecular complex with TRAP, PAT and possibly other components in sporozoites remains to be established. Future studies will also determine how precisely CLAMP regulates TRAP shedding. CLAMP could participate in the secretion of TRAP, and possibly other adhesins, from the sporozoite micronemes. Alternatively, it might regulate the proteolytic processing of TRAP at the sporozoite surface following microneme exocytosis (Ejigiri et al., 2012). Interestingly, in *Toxoplasma* tachyzoites, depletion of CLAMP does not impact the secretion and shedding of MIC2, a micronemal adhesin related to TRAP (Sidik et al., 2016), suggesting that the protein may have different functions in *Toxoplasma* and *Plasmodium* parasites.

CLAMP may also participate in the gliding motility of merozoites, which has recently emerged as an important precursor step during invasion of erythrocytes (Yahata et al., 2021). While TRAP is not expressed in merozoites and, unlike CLAMP, is dispensable for *Plasmodium* blood stage growth, merozoites express two members of the TRAP protein family, the merozoite thrombospondin-related anonymous protein (MTRAP) and the *Plasmodium* thrombospondin-related apical membrane protein (PTRAMP) (Baum et al., 2006; Morahan et al., 2009; Thompson et al., 2004). Unlike MTRAP, which is required in sexual blood stages for gamete egress (Bargieri et al., 2016; Kehrer et al., 2016a), PTRAMP is essential during asexual blood stage growth in both *P. falciparum* and *P. berghei* (Bushell et al., 2017; Zhang et al., 2018). Whether CLAMP participates in merozoite gliding motility and/or regulates the function of merozoite adhesins, such as PTRAMP, deserves further investigation.

In addition to TRAP, various proteins have been identified as involved in sporozoite gliding motility, such as the TRAP-related protein (TREP) (Combe et al., 2009; Steinbuechel and Matuschewski, 2009), LIMP (Santos et al., 2017) or the Secreted Protein with Altered Thrombospondin Repeat (SPATR) (Costa et al., 2022). Like CLAMP, SPATR is conserved in Apicomplexa and is essential in asexual blood stages of the parasite life cycle. Interestingly, a recent study based on conditional knockdown in *P. berghei* showed that sporozoites lacking SPATR have impaired motility, strongly reduced capacity to invade salivary glands and decreased infectivity to mice (Costa et al., 2022), a phenotype that is similar to the one we observed with CLAMP-deficient parasites. Costa *et al*. excluded a role of SPATR in TRAP secretion, based on the detection of TRAP at the surface of mutant sporozoites by immunofluorescence (Costa et al., 2022). In our hands, surface staining of TRAP did not show any overt difference between untreated and rapamycin-treated parasites, irrespective of stimulation of microneme secretion (**S3 Fig**). It is possible however that surface immunostaining may not capture the entire dynamics of TRAP secretion and shedding.

In summary, we show here that CLAMP plays an essential role across invasive stages of the malaria parasite, and is required for gliding motility and infectivity of *P. berghei* sporozoites by regulating the secretion of TRAP. CLAMP may thus represent a potential new target for antimalarial strategies. In this regard, a recent study showed that CLAMP-specific antibodies can inhibit invasion of equine erythrocytes by the apicomplexan *Theileria equi*, and identified neutralization-sensitive epitopes in the predicted extracellular loops of the protein (Onzere et al., 2021). It will be important to test whether *Plasmodium* CLAMP can be similarly targeted by neutralizing antibodies. This would open interesting perspectives for interventions acting both against merozoites, to prevent invasion of erythrocytes, and sporozoites, to block infection early after transmission by the mosquito.

## Materials and methods

### Ethics statement

All animal work was conducted in strict accordance with the Directive 2010/63/EU of the European Parliament and Council on the protection of animals used for scientific purposes. Protocols were approved by the Ethical Committee Charles Darwin N°005 (approval #7475-2016110315516522).

### Experimental animals, parasites and cell lines

Female Swiss mice (6–8 weeks old, from Janvier Labs) were used for all routine parasite infections. Parasite transfections were performed in the parental PbDiCre line (ANKA strain), which constitutively express the DiCre components in addition to a mCherry fluorescence cassette (Fernandes et al., 2020). Parasite infections in mice were initiated through intraperitoneal injections of infected RBCs. A drop of blood was collected from the tail in 1ml PBS daily and used to monitor the parasitaemia by flow cytometry. *Anopheles stephensi* mosquitoes were reared at 24°C with 80 % humidity and permitted to feed on anaesthetised infected mice, using standard methods of mosquito infection as previously described (Ramakrishnan et al., 2013). Post-feeding, *P. berghei*-infected mosquitoes were kept at 21°C and fed on a 10% sucrose solution. Midgut and haemolymph sporozoites were collected at day 16 and salivary gland sporozoites at day 21 post-infection. Sporozoites were collected by hand dissection and homogenisation of isolated midguts or salivary glands in complete DMEM medium (supplemented with 10% FBS, 1% Penicillin-Streptomycin and 1% L-Glutamine), then counted in a Neubauer haemocytometer. HepG2 cells (ATCC HB-8065) were cultured in complete DMEM in collagen-coated plates, at 37°C, 5% CO2, as previously described (Silvie et al., 2007), and used for invasion and cell traversal assays.

### Generation of the *clamp*cKO parasite line in *P. berghei*

#### Plasmid constructs

Two plasmids, P1 and P2, were generated for insertion of LoxN sites upstream and downstream of *clamp*, respectively. The P1 plasmid was assembled by inserting two homologous sequences, 5’HR1 and 5’HR2, both localized upstream of *clamp* promoter region, in the pUpstream2Lox plasmid (Addgene #164573), which contains a GFP-2A-hDHFR cassette flanked by two LoxN sites. Both 5’HR1 (895 bp) and 5’HR2 (1014 bp) were amplified by PCR from WT PbANKA genomic DNA, and inserted into *Kpn*I/*Xho*I and *Nhe*I sites, respectively, of the pUpstream2Lox plasmid. The P2 plasmid was assembled by inserting two homologous sequences, 3’HR1 and 3’HR2, corresponding to the end of *clamp* ORF and to *clamp* 3’UTR, respectively, in the pDownstream1Lox plasmid (Addgene #164574), which contains a GFP-2A-hDHFR cassette followed by a single LoxN site. Both 3’HR1 (862 bp) and 3’HR2 (999 bp) were amplified by PCR from WT PbANKA genomic DNA, and inserted into *Kpn*I/*Xho*I and *Nhe*I sites, respectively, of the pDownstream1Lox plasmid. A triple Flag epitope tag was inserted in frame with *clamp* ORF immediately before the stop codon. In addition, a 559 bp fragment corresponding to the 3’ UTR sequence from *P. yoelii clamp* gene (PY17X_0515300) was inserted immediately downstream of 3’HR1, to allow proper gene expression and avoid spontaneous recombination with the 3’ UTR of *P. berghei* clamp, which was used as 3’HR2. Plasmids P1 and P2 were verified by Sanger DNA sequencing (Eurofins Genomics) and linearized with *Kpn*I and *Nhe*I before transfection. All the primers used for plasmid assembly are listed in **S2 Table**.

#### Transfection and selection

Parental DiCre parasites were transfected with the P1 plasmid to generate *clamp*-P1 parasites, which were then exposed to rapamycin and transfected with the P2 plasmid to generate the final *clamp*cKO line. For the first transfection, schizonts purified from an overnight culture of PbDiCre blood stage parasites were transfected with 10 µg of linearized P1 plasmid using the AMAXA Nucleofector device (program U033), as previously described (Janse et al., 2006), and immediately injected intravenously into the tail vein of SWISS mice. To permit the selection of resistant transgenic parasites, pyrimethamine (35 mg/L) and 5-flurocytosine (0.5 mg/ml) were added to the mouse drinking water, starting one day after transfection. The parasitaemia was monitored daily by flow cytometry and the mice sacrificed at a parasitaemia of 2-3%, allowing preparation of frozen parasite stocks and isolation of parasites for genomic DNA extraction. After drug selection, GFP^+^/mCherry^+^ *clamp*-P1 parasites were sorted by flow cytometry. Mice were injected intraperitoneally with frozen parasite stocks and monitored until the parasitaemia was between 0.1 and 1%. On the day of sorting, one drop of tail blood was collected in 1ml PBS and used for sorting of 100 iRBCs on a FACSAria II (Becton-Dickinson), as described (Manzoni et al., 2014). Sorted parasites were recovered in 200 µl RPMI containing 20% FBS and injected intravenously into two mice. *clamp*-P1 parasites were exposed to a single dose of rapamycin to induce excision of the GFP-2A-hDHFR cassette. The resulting GFP^-^/mCherry^+^ *clamp*-P1 parasites, which retained a single LoxN site inserted upstream of *clamp*, were sorted by flow cytometry and amplified in mice. Rapamycin-exposed *clamp*-P1 parasites were then transfected with the P2 plasmid, as described above, to generate the *clamp*cKO line. GFP^+^/mCherry^+^ *clamp*cKO parasites were sorted by flow cytometry, as described above, and cloned by limiting dilution and injection into mice to generate the final population used in this study. Parasites were genotyped at each step to verify the correct integration of the constructs and the absence of non-recombined *clamp* locus.

### Genotyping PCR

The blood collected from infected mice was passed through a CF11 column (Whatman) to deplete leucocytes. The collected RBCs were then centrifuged and lysed with 0.2% saponin (Sigma), before genomic DNA isolation using the DNA Easy Blood and Tissue Kit (Qiagen), according to the manufacturer’s instructions. Specific PCR primers were designed to check for wild-type and recombined loci and are listed in **S2 Table**. PCR reactions were carried out using Recombinant Taq DNA Polymerase (Thermo Scientific) and standard PCR cycling conditions.

### Rapamycin-induced gene excision

*clamp*cKO-infected mice were administered a single dose of 200 μg rapamycin (Rapamune^®^, Pfizer) by oral gavage. Treatment efficacy was validated by the observation of a decrease in GFP fluorescence intensity in circulating blood-stage parasites after treatment, reflecting gene excision. Rapamycin-treated *clamp*cKO parasites were transmitted to mosquitoes one day after rapamycin administration to mice, as described (Fernandes et al., 2021, 2020).

#### *In vitro* infection assays

HepG2 cells cultured in DMEM complete medium were seeded at a density of 30,000 cells/well in a 96-well plate for flow cytometry analysis or 100,000 cells/well in 96-well µ-slide (Ibidi) for immunofluorescence assays, 24 hours prior to addition of sporozoites. Culture medium was refreshed with complete DMEM on the day of infection, followed by incubation with 3,000 or 1,000 sporozoites, for flow cytometry or immunofluorescence assays, respectively. For quantification of traversal events, fluorescein-conjugated dextran (0.5 mg/ml, Life Technologies) was added to the wells together with sporozoites. After 3 hours, cells were washed twice with PBS, trypsinized, then resuspended in complete DMEM for analysis by flow cytometry on a Guava EasyCyte 6/2L bench cytometer equipped with 488nm and 532nm lasers (Millipore). Control wells were prepared without sporozoites to measure the basal level of dextran uptake. To quantify parasite liver stage infection, cells were washed twice with complete DMEM 3 hours after sporozoite addition, and then incubated for another 24h. Cultures were then fixed with 4% PFA, followed by two washes with PBS. Cells were then quenched with 0.1M glycine for 5 min, washed twice with PBS then permeabilized with 1% Triton X-100 for 5 min before 2 washes in PBS and blocking in PBS + 3% BSA. Samples were then stained with goat anti-UIS4 primary antibodies (1:500, Sicgen), followed by donkey anti-goat Alexa Fluor 594 secondary antibodies (1:1000, Life Technologies), both diluted in PBS with 3% BSA. EEFs were then counted based on the presence of a UIS4-stained PV.

### Imaging of parasites

Midguts and salivary glands were carefully collected from infected mosquitoes, mounted in PBS and directly examined without fixation on a fluorescence microscope. Sporozoites were isolated by disruption of midguts or salivary glands and resuspended in PBS, placed on a coverslip then fixed for 10 min with 4% PFA followed by two washes with PBS. For immunostaining, sporozoites were fixed and permeabilized, and incubated with 3% BSA in PBS for 1h. Sporozoites were then stained with anti-Flag primary mouse antibody (M2 clone, Sigma), washed twice with PBS then incubated with Alexa Fluor anti-mouse 647 secondary antibodies (Life Technologies) and Hoechst 77742 (Life Technologies), before examination by fluorescence microscopy. All images were acquired on an Axio Observer Z1 Zeiss fluorescence microscope using the Zen software (Zeiss). The same exposure time was set for rapamycin-exposed and untreated *clamp*cKO parasites in order to allow comparisons. Images were processed with ImageJ for adjustment of contrast.

### Motility assay

For motility assays, rapamycin-exposed and untreated *clamp*cKO sporozoites were collected from manual dissection of infected mosquitoes and mixed at a 1:1 ratio, and kept in PBS on ice until imaging in PBS with 3% BSA. Sporozoites were then deposited in BSA-coated 96-well plates, centrifuged for 2 min at 100 x g and placed at 37°C in the microscope chamber with 5% CO2. Acquisitions were initiated after 10 min, to allow sporozoite activation, with one image captured every second for a total of 3 min. Images were acquired on an Axio Observer 7 widefield microscope (Zeiss), with ORCA-Fusion Digital CMOS cameras and a Duolink camera adapter for simultaneous two-color acquisition with integrated multi-bandpass emission filter cubes for efficient image acquisition. The microscope was equipped with a 20x objective and ZEN acquisition software. Movies (181 frames per movie) were projected into a single image for each channel and merged using ImageJ. Sporozoites were counted manually based on their movement pattern and divided into four groups: circular gliding, floating, waving or fixed. Rapamycin-exposed and untreated *clamp*cKO sporozoites were differentiated based on their fluorescence (after merge: red for CLAMP-deficient rapamycin-exposed parasites and yellow for control untreated parasites). Three independent motility experiments were performed, with a total of 1496 and 864 sporozoites analyzed in the untreated and rapamycin-treated conditions, respectively.

### Immunoprecipitation assay and mass spectrometry analysis

Freshly dissected untreated *clamp*cKO sporozoites were lysed on ice for 30 min in a lysis buffer containing 0.5% w/v NP40 (Igepal CA-630, Sigma) and protease inhibitors. After centrifugation (15,000 × g, 15 min, 4°C), supernatants were collected and incubated with protein G-conjugated sepharose for preclearing overnight. Precleared lysates were subjected to CLAMP-Flag immunoprecipitation using Anti-FLAG M2 Affinity Gel (Sigma) for 2h at 4°C, according to the manufacturer’s protocol. PbGFP parasites with untagged proteins were used as a control and treated in the same fashion. After washes, proteins on beads were eluted in 2X Laemmli and denatured (95°C, 5 min). After centrifugation, supernatants were collected for further analysis. Samples were subjected to a short SDS-PAGE migration, and gel pieces were processed for protein trypsin digestion by the DigestProMSi robot (Intavis), as described (Hamada et al., 2021). Peptide samples were analyzed on a timsTOF PRO mass spectrometer (Bruker) coupled to the nanoElute HPLC, as described (Hamada et al., 2021). Mascot generic files were processed with X!Tandem pipeline (version 0.2.36) using the PlasmoDB_PB_39_PbergheiANKA database, as described (Hamada et al., 2021). Data were obtained from 2 independent *clamp*cKO (untreated) and 3 control PbGFP sporozoite lysates, and are provided in **S1 Table**.

### Western-blot analysis

Rapamycin-exposed and untreated *clamp*cKO sporozoites were isolated from the haemolymph of infected mosquitoes at day 16 post-transmission and resuspended in 1X PBS. For analysis of TRAP shedding, microneme secretion was stimulated by incubation for 15 min at 37°C in a buffer containing 1% BSA and 1% ethanol, as described (Brown et al., 2020). Pellet and supernatant fractions were then isolated by centrifugation, resuspended in Laemmli buffer and analyzed by SDS-PAGE under non-reducing conditions. Western blotting was performed using rabbit polyclonal antibodies against TRAP (Klug et al., 2020), 3D11 monoclonal antibody against CSP (Yoshida et al., 1980), or M2 monoclonal antibody against Flag (Sigma), and secondary antibodies coupled with Alexa Fluor 680 or 800. Membranes were then analyzed using the InfraRed Odyssey system (Licor). Band intensities were quantified using ImageJ.

### Statistical Analysis

Statistical significance was assessed by two-way ANOVA, ratio paired t tests or Chi-squared test, as indicated in the figure legends. All statistical tests were computed with GraphPad Prism 7 (GraphPad Software). *In vitro* experiments were performed with a minimum of three technical replicates per experiment. Quantitative source data are provided in **S3 Table**.

## Acknowledgements

We thank Maurel Tefit and Thierry Houpert for rearing of mosquitoes, and Freddy Frischknecht and Jessica Kehrer for the kind gift of TRAP antibodies. This work was funded by grants from the Laboratoire d’Excellence ParaFrap (ANR-11-LABX-0024), the Agence Nationale de la Recherche (ANR-20-CE18-0013) and the Fondation pour la Recherche Médicale (EQU201903007823). The authors acknowledge the Conseil Régional d’Ile-de-France, Sorbonne Université, the National Institute for Health and Medical Research (INSERM) and the Biology, Health and Agronomy Infrastructure (IBiSA) for funding the timsTOF PRO. This work benefited from equipment and services from the ICM.Quant core facility (Paris Brain Institute), a platform supported through the ANR grants, ANR-10-IAIHU-06 and ANR-11-INBS-0011-NeurATRIS. ML was supported by a ‘DIM 1Health’ doctoral fellowship awarded by the Conseil Régional d’Ile-de-France.

## Supporting information

### Supplemental tables

**S1 Table**. *P. berghei* proteins identified by mass spectrometry after anti-Flag immunoprecipitation.

**S2 Table**. List of oligonucleotides used in the study.

**S3 Table**. Quantitative source data and statistical analysis.

### Supplemental figures

**S1 Fig.**
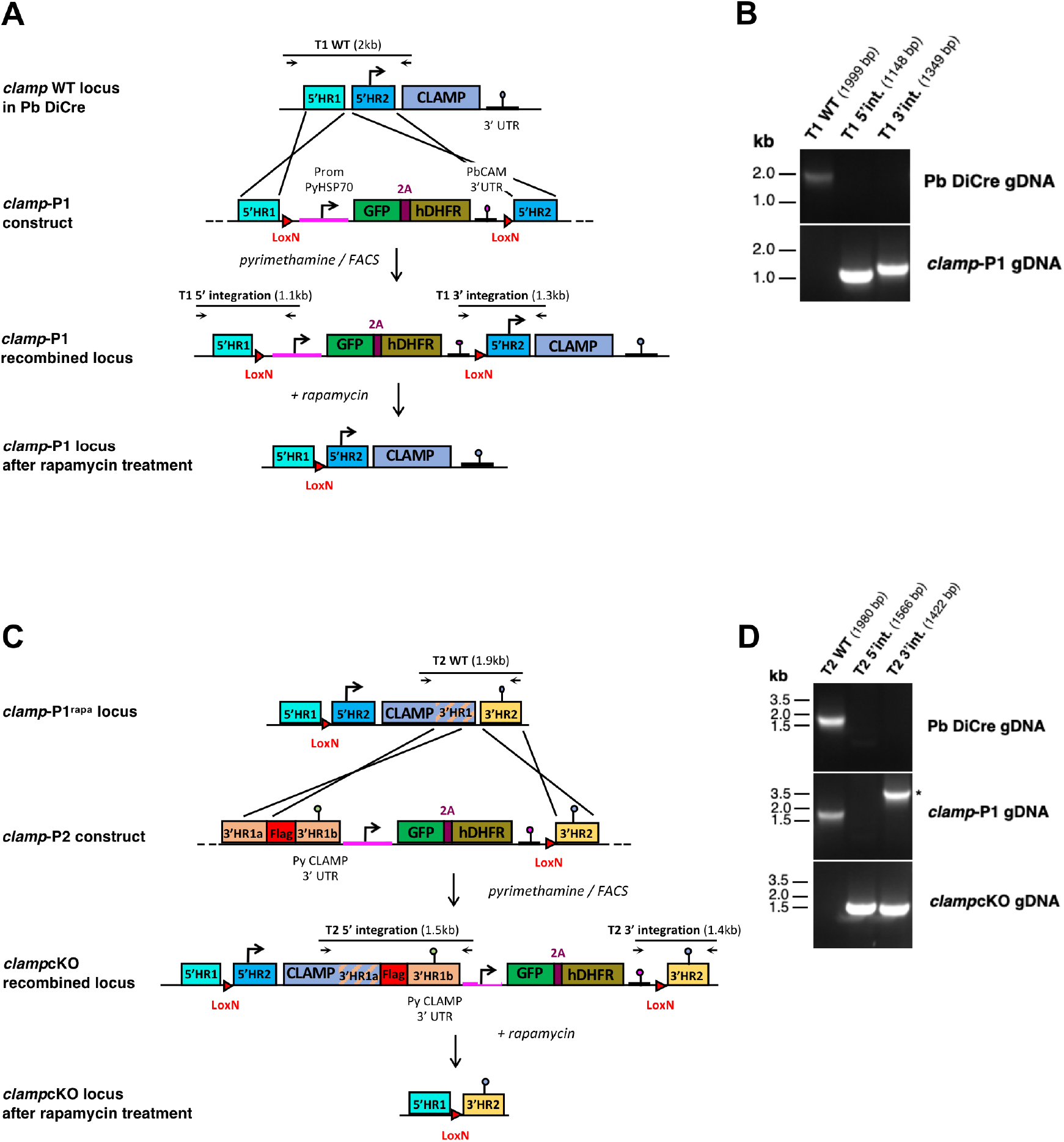
Generation of *P. berghei clamp*cKO parasites. **A**. Detailed strategy to insert a LoxN site upstream of *clamp* gene using the P1 construct. Upstream homology regions (5’HR1 and 5’HR2) were inserted in the pUpstream2Lox plasmid on each side of a GFP-2A-hDHFR cassette flanked by two LoxN sites. The P1 construct was transfected into mCherry-expressing PbDiCre parasites. Following parasite transfection and selection with pyrimethamine, mCherry^+^/GFP^+^ parasites were sorted by flow cytometry to exclude any residual GFP^-^ population. Rapamycin-induced excision lead to removal of the GFP-2A-hDHFR cassette and the retention of a single LoxN site upstream of *clamp*. Genotyping primers and expected PCR fragments are indicated by arrows and lines, respectively. **B**. PCR analysis of genomic DNA isolated from parental PbDiCre and *clamp*-P1 parasites. Confirmation of the predicted recombination events was assessed with primer combinations specific for WT, 5’ or 3’ integration for the first transfection (T1). Primers used for genotyping are listed in **S2 Table. C**. Detailed strategy to insert a LoxN site downstream of *clamp* gene using the P2 construct. Downstream 3’ homology regions (3’HR1 and 3’HR2) were inserted in the pDownstream1Lox plasmid on each side of a GFP-2A-hDHFR cassette, flanked on one side by a single LoxN site. A triple Flag epitope tag (3xFlag) was inserted in frame with *clamp* ORF immediately before the STOP codon. In addition, a 559 bp fragment corresponding to the 3’ UTR sequence from *P. yoelii clamp* gene was inserted immediately downstream of STOP codon, to allow proper gene expression and avoid spontaneous recombination with the 3’ UTR of *P. berghei* clamp, which was used as 3’HR2. The P2 construct was transfected into rapamycin-treated mCherry^+^/GFP^-^ *clamp*-P1 parasites (*clamp*-P1^rapa^). Following parasite transfection and selection with pyrimethamine, mCherry^+^/GFP^+^ parasites were sorted by flow cytometry to exclude any residual GFP^-^ population, and cloned by limiting dilution and injection into mice, resulting in the final *clamp*cKO parasite line. Exposure of *clamp*cKO parasites to rapamycin leads to excision of *clamp* gene together with the GFP-2A-hDHFR cassette. Genotyping primers and expected PCR fragments are indicated by arrows and lines, respectively. **D**. PCR analysis of genomic DNA isolated from parental PbDiCre, *clamp*-P1 and *clamp*cKO parasites. Confirmation of predicted recombination events was assessed with primer combinations specific for WT, 5’ or 3’ integration for the second transfection (T2). A band in *clamp*-P1 gDNA for 3’ integration is due to the forward primer which can bind to the PbCAM 3’ UTR sequence integrated upstream of *clamp* during the first transfection, and is indicated by an asterisk. Primers used for genotyping are listed in **S2 Table**.

**S2 Fig.**
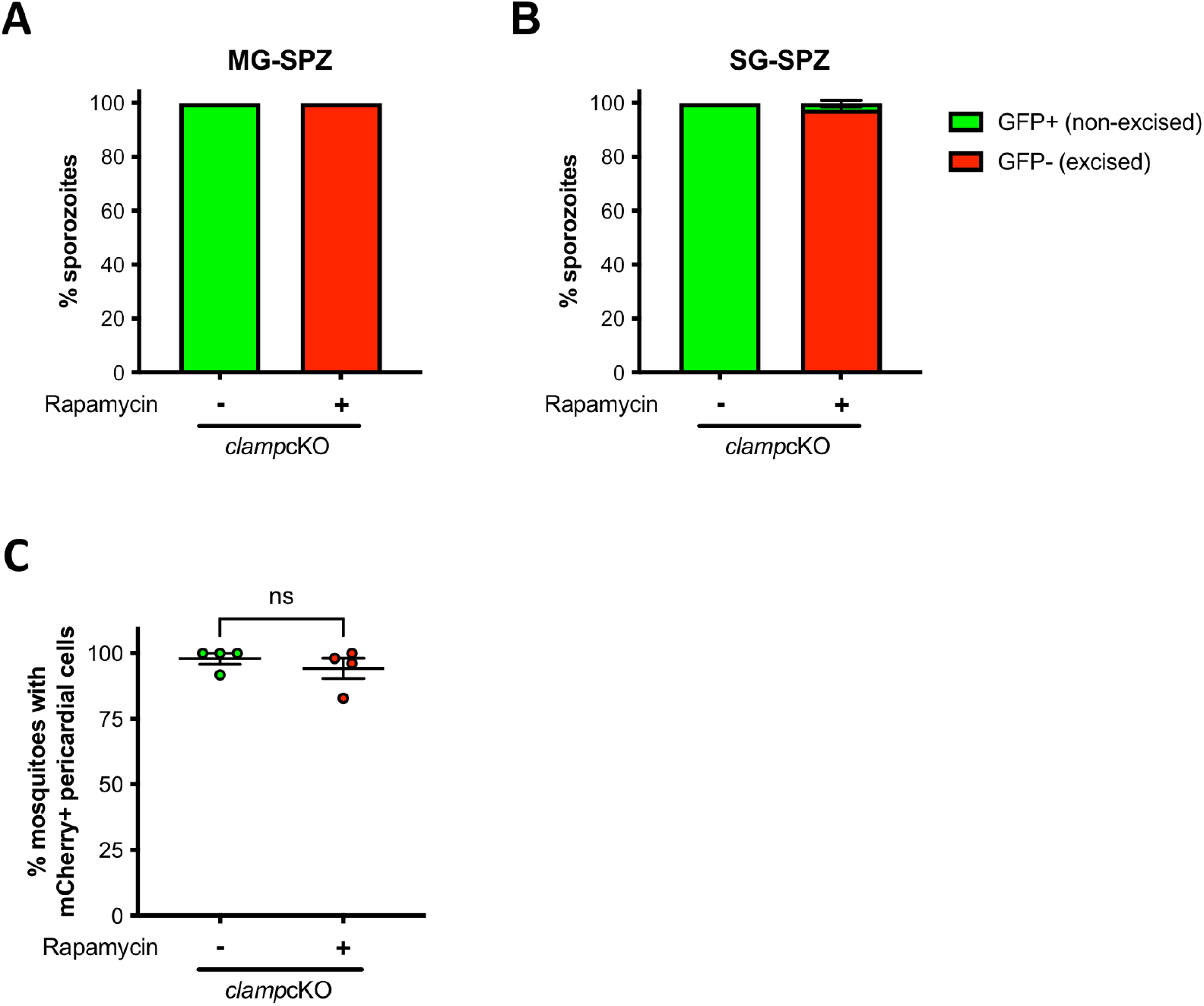
Analysis of *clamp*cKO parasite development in mosquitoes. **A-B**. Fluorescence-based quantification of excised (mCherry^+^GFP^-^) and non-excised (mCherry^+^GFP^+^) sporozoites isolated from midguts (A) and salivary glands (B) of female mosquitoes infected with rapamycin-exposed and untreated *clamp*cKO parasites. Results shown are based on observation of at least 200 sporozoites per condition and per experiment (mean +/-SEM of four independent experiments). **C**. Quantification of infected female mosquitoes exhibiting mCherry-labelled pericardial cells 16 days post-infection, based on observation of at least 50 mosquitoes per condition (mean +/-SEM of four independent experiments). Ns, non-significant (Two-tailed ratio paired t test).

**S3 Fig.**
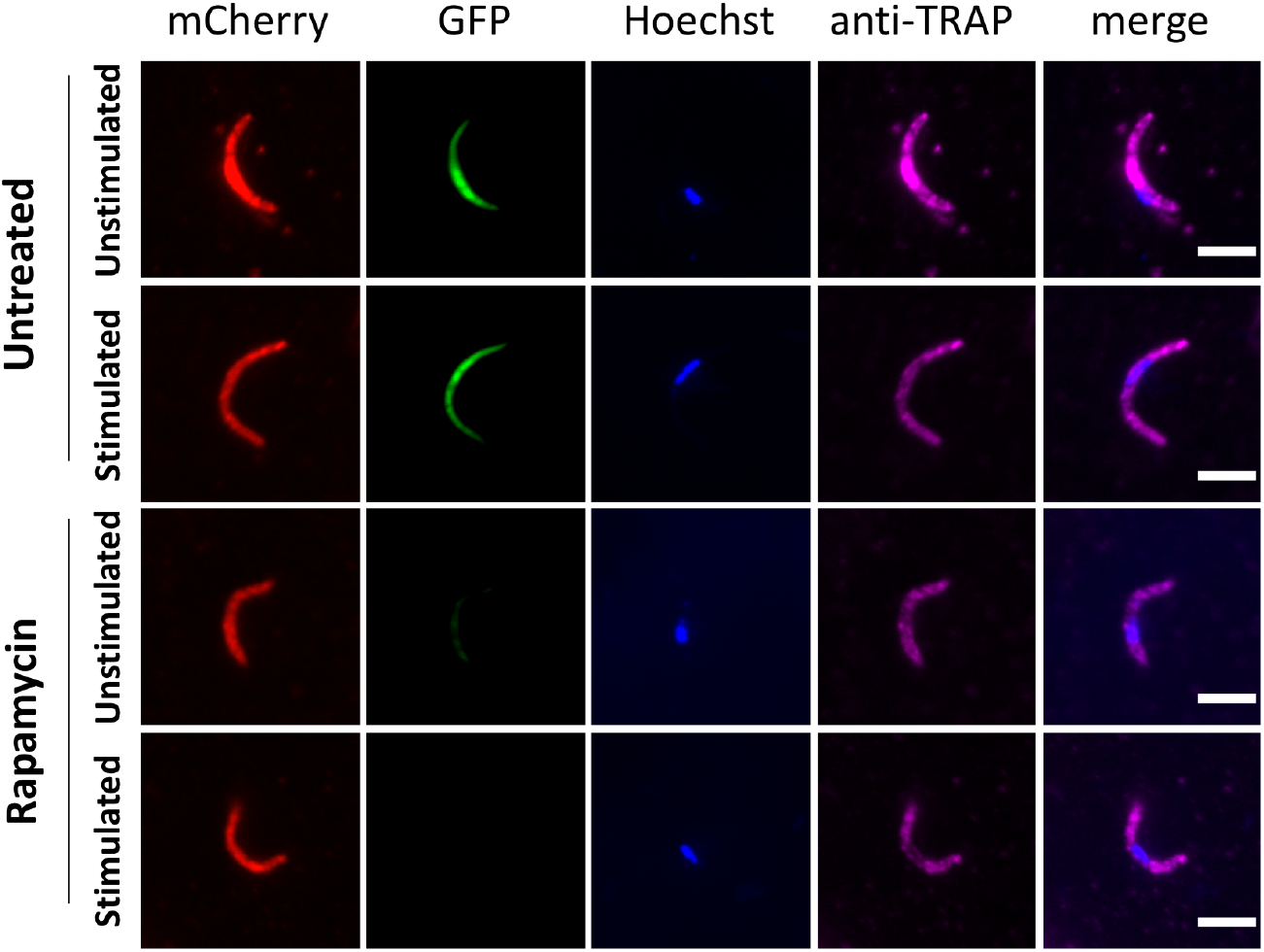
TRAP expression on the surface of salivary gland sporozoites. Sporozoites were isolated from the salivary glands of mosquitoes infected with untreated or rapamycin-exposed *clamp*cKO parasites. Microneme secretion was stimulated by incubation at 37°C in the presence of 1% BSA and 1% ethanol for 15 min. Stimulated and unstimulated sporozoites were then fixed with 4% PFA without permeabilization, and stained with anti-TRAP antibodies (magenta) and the nuclear stain Hoechst 33342 (blue). Untreated parasites express GFP (green) and mCherry (red), while rapamycin-treated parasites express mCherry only. Scale bars, 5 μm.

### Supplemental movies

**Movie 1. Motility of untreated and rapamycin-exposed *clamp*cKO salivary gland sporozoites**. Untreated (mCherry^+^/GFP^+^, yellow) and rapamycin-treated (mCherry^+^/GFP^-^, red) *clamp*cKO sporozoites were mixed at a 1:1 ratio. Motility was recorded with one frame per second (fps) for 3 min after activation in 3% BSA. Video was edited with 7fps. Only untreated parasites show circular gliding activity.

## References

Aliprandini E, Tavares J, Panatieri RH, Thiberge S, Yamamoto MM, Silvie O, Ishino T, Yuda M, Dartevelle S, Traincard F, Boscardin SB, Amino R. 2018. Cytotoxic anti-circumsporozoite antibodies target malaria sporozoites in the host skin. Nat Microbiol. doi:10.1038/s41564-018-0254-z

Andenmatten N, Egarter S, Jackson AJ, Jullien N, Herman J-P, Meissner M. 2012. Conditional genome engineering in Toxoplasma gondii uncovers alternative invasion mechanisms. Nat Methods 10:125–127. doi:10.1038/nmeth.2301

Arredondo SA, Swearingen KE, Martinson T, Steel R, Dankwa DA, Harupa A, Camargo N, Betz W, Vigdorovich V, Oliver BG, Kangwanrangsan N, Ishino T, Sather N, Mikolajczak S, Vaughan AM, Torii M, Moritz RL, Kappe SHII. 2018. The Micronemal Plasmodium Proteins P36 and P52 Act in Concert to Establish the Replication-Permissive Compartment Within Infected Hepatocytes 8:413. doi:10.3389/fcimb.2018.00413

Bargieri D, Lagal V, Andenmatten N, Tardieux I, Meissner M, Ménard R. 2014. Host Cell Invasion by Apicomplexan Parasites: The Junction Conundrum. PLoS Pathog 10:1–9. doi:10.1371/journal.ppat.1004273

Bargieri DY, Thiberge S, Tay CL, Carey AF, Rantz A, Hischen F, Lorthiois A, Straschil U, Singh P, Singh S, Triglia T, Tsuboi T, Cowman A, Chitnis C, Alano P, Baum J, Pradel G, Lavazec C, Ménard R. 2016. Plasmodium Merozoite TRAP Family Protein Is Essential for Vacuole Membrane Disruption and Gamete Egress from Erythrocytes. Cell Host Microbe 20:618–630. doi:10.1016/j.chom.2016.10.015

Baum J, Richard D, Healer J, Rug M, Krnajski Z, Gilberger TW, Green JL, Holder AA, Cowman AF. 2006. A conserved molecular motor drives cell invasion and gliding motility across malaria life cycle stages and other apicomplexan parasites. J Biol Chem 281. doi:10.1074/jbc.M509807200

Besteiro S, Dubremetz JF, Lebrun M. 2011. The moving junction of apicomplexan parasites: A key structure for invasion. Cell Microbiol 13:797–805. doi:10.1111/j.1462-5822.2011.01597.x

Brown KM, Sibley LD, Lourido S. 2020. High-Throughput Measurement of Microneme Secretion in Toxoplasma gondiiMethods in Molecular Biology. doi:10.1007/978-1-4939-9857-9_9

Bushell E, Gomes AR, Sanderson T, Anar B, Girling G, Herd C, Metcalf T, Modrzynska K, Schwach F, Martin RE, Mather MW, McFadden GI, Parts L, Rutledge GG, Vaidya AB, Wengelnik K, Rayner JC, Billker O. 2017. Functional Profiling of a Plasmodium Genome Reveals an Abundance of Essential Genes. Cell 170:260-272.e8. doi:10.1016/j.cell.2017.06.030

Combe A, Moreira C, Ackerman S, Thiberge S, Templeton TJ, Ménard R. 2009. TREP, a novel protein necessary for gliding motility of the malaria sporozoite. Int J Parasitol 39. doi:10.1016/j.ijpara.2008.10.004

Costa DM, Sá M, Teixeira AR, Pérez-Cabezas B, Golba S, Sefiane-Djemaoune H, Formaglio P, Franke-Fayard B, Janse CJ, Amino R, Tavares J. 2022. Downregulation of the secreted protein with an altered thrombospondin repeat (SPATR) impacts the infectivity of malaria sporozoites. bioRxiv.

Cowman AF, Tonkin CJ, Tham W-H, Duraisingh MT. 2017. The Molecular Basis of Erythrocyte Invasion by Malaria Parasites. Cell Host Microbe 22:232–245. doi:10.1016/j.chom.2017.07.003

Ejigiri I, Ragheb DRT, Pino P, Coppi A, Bennett BL, Soldati-Favre D, Sinnis P. 2012. Shedding of TRAP by a rhomboid protease from the malaria sporozoite surface is essential for gliding motility and sporozoite infectivity. PLoS Pathog 8:e1002725. doi:10.1371/journal.ppat.1002725

Fernandes P, Briquet S, Patarot D, Loubens M, Hoareau-Coudert B, Silvie O. 2020. The dimerisable Cre recombinase allows conditional genome editing in the mosquito stages of Plasmodium berghei. PLoS One 15. doi:10.1371/journal.pone.0236616

Fernandes P, Loubens M, Le Borgne R, Marinach C, Ardin B, Briquet S, Vincensini L, Hamada S, Hoareau-Coudert B, Verbavatz J-M, Weiner A, Silvie O. 2022. The AMA1-RON complex drives <I>Plasmodium</I> sporozoite invasion in the mosquito and mammalian hosts. bioRxiv 2022.01.04.474787. doi:10.1101/2022.01.04.474787

Fernandes P, Loubens M, Silvie O, Briquet S. 2021. Conditional Gene Deletion in Mammalian and Mosquito Stages of Plasmodium berghei Using Dimerizable Cre RecombinaseMethods in Molecular Biology. doi:10.1007/978-1-0716-1681-9_7

Frénal K, Dubremetz J-F, Lebrun M, Soldati-Favre D. 2017. Gliding motility powers invasion and egress in Apicomplexa. Nat Rev Microbiol 15:645–660. doi:10.1038/nrmicro.2017.86

Hamada S, Pionneau C, Parizot C, Silvie O, Chardonnet S, Marinach C. 2021. In-depth proteomic analysis of Plasmodium berghei sporozoites using trapped ion mobility spectrometry with parallel accumulation-serial fragmentation. Proteomics 21. doi:10.1002/pmic.202000305

Hart RJ, Lawres L, Fritzen E, Mamoun C Ben, Aly ASI. 2014. Plasmodium yoelii vitamin B5 pantothenate transporter candidate is essential for parasite transmission to the mosquito. Sci Rep 4. doi:10.1038/srep05665

Ishino T, Chinzei Y, Yuda M. 2005a. A Plasmodium sporozoite protein with a membrane attack complex domain is required for breaching the liver sinusoidal cell layer prior to hepatocyte infection. Cell Microbiol 7:199–208. doi:10.1111/j.1462-5822.2004.00447.x

Ishino T, Chinzei Y, Yuda M. 2005b. Two proteins with 6-cys motifs are required for malarial parasites to commit to infection of the hepatocyte. Mol Microbiol 58:1264–1275. doi:10.1111/j.1365-2958.2005.04801.x

Ishino T, Murata E, Tokunaga N, Baba M, Tachibana M, Thongkukiatkul A, Tsuboi T, Torii M. 2019. Rhoptry neck protein 2 expressed in Plasmodium sporozoites plays a crucial role during invasion of mosquito salivary glands. Cell Microbiol 21:e12964. doi:10.1111/cmi.12964

Ishino T, Yano K, Chinzei Y, Yuda M. 2004. Cell-passage activity is required for the malarial parasite to cross the liver sinusoidal cell layer. PLoS Biol 2:77–84. doi:10.1371/journal.pbio.0020004

Janse CJ, Ramesar J, Waters AP. 2006. High-efficiency transfection and drug selection of genetically transformed blood stages of the rodent malaria parasite Plasmodium berghei. Nat Protoc 1:346–356. doi:10.1038/nprot.2006.53

Jullien N, Sampieri F, Enjalbert A, Herman JP. 2003. Regulation of Cre recombinase by ligand-induced complementation of inactive fragments. Nucleic Acids Res 31. doi:10.1093/nar/gng131

Kappe S, Bruderer T, Gantt S, Fujioka H, Nussenzweig V, Ménard R. 1999. Conservation of a gliding motility and cell invasion machinery in Apicomplexan parasites. J Cell Biol 147. doi:10.1083/jcb.147.5.937

Kehrer J, Frischknecht F, Mair GR. 2016a. Proteomic analysis of the plasmodium berghei gametocyte egressome and vesicular bioid of osmiophilic body proteins identifies merozoite trap-like protein (MTRAP) as an essential factor for parasite transmission. Mol Cell Proteomics 15. doi:10.1074/mcp.M116.058263

Kehrer J, Singer M, Lemgruber L, Silva PAGC, Frischknecht F, Mair GR. 2016b. A Putative Small Solute Transporter Is Responsible for the Secretion of G377 and TRAP-Containing Secretory Vesicles during Plasmodium Gamete Egress and Sporozoite Motility. PLoS Pathog 12. doi:10.1371/journal.ppat.1005734

Kenthirapalan S, Waters AP, Matuschewski K, Kooij TWA. 2016. Functional profiles of orphan membrane transporters in the life cycle of the malaria parasite. Nat Commun 7. doi:10.1038/ncomms10519

Klug D, Goellner S, Kehrer J, Sattler J, Strauss L, Singer M, Lu C, Springer TA, Frischknecht F. 2020. Evolutionarily distant I domains can functionally replace the essential ligandbinding domain of plasmodium trap. Elife 9:1–27. doi:10.7554/eLife.57572

Lindner SE, Swearingen KE, Harupa A, Vaughan AM, Sinnis P, Moritz RL, Kappe SH. 2013. Total and putative surface proteomics of malaria parasite salivary gland sporozoites. Mol Cell Proteomics 12:1127–1143. doi:10.1074/mcp.M112.024505

Manzoni G, Briquet S, Risco-Castillo V. 2014. A rapid and robust selection procedure for generating drug-selectable marker-free recombinant malaria parasites. Sci Rep 99210:1–10. doi:10.1038/srep04760

Manzoni G, Marinach C, Topçu S, Briquet S, Grand M, Tolle M, Gransagne M, Lescar J, Andolina C, Franetich JF, Zeisel MB, Huby T, Rubinstein E, Snounou G, Mazier D, Nosten F, Baumert TF, Silvie O. 2017. Plasmodium P36 determines host cell receptor usage during sporozoite invasion. Elife 6:e25903. doi:10.7554/eLife.25903

Matuschewski K, Nunes AC, Nussenzweig V, Menard R. 2002. Plasmodium sporozoite invasion into insect and mammalian cells is directed by the same dual binding system. Embo J 21:1597–1606.

Morahan BJ, Wang L, Coppel RL. 2009. No TRAP, no invasion. Trends Parasitol. doi:10.1016/j.pt.2008.11.004

Mota MM, Pradel G, Vanderberg JP, Hafalla JC, Frevert U, Nussenzweig RS, Nussenzweig V, Rodriguez A. 2001. Migration of Plasmodium sporozoites through cells before infection. Science 291:141–144.

Mueller AK, Camargo N, Kaiser K, Andorfer C, Frevert U, Matuschewski K, Kappe SHI. 2005. Plasmodium liver stage developmental arrest by depletion of a protein at the parasite-host interface. Proc Natl Acad Sci U S A 102:3022–3027. doi:10.1073/pnas.0408442102

Nozaki M, Baba M, Tachibana M, Tokunaga N, Torii M, Ishino T. 2020. Detection of the Rhoptry Neck Protein Complex in Plasmodium Sporozoites and Its Contribution to Sporozoite Invasion of Salivary Glands. mSphere 5:e00325–20. doi:10.1128/msphere.00325-20

Onzere CK, Fry LM, Bishop RP, Silva MG, Bastos RG, Knowles DP, Suarez CE. 2021. Theileria equi claudin like apicomplexan microneme protein contains neutralization-sensitive epitopes and interacts with components of the equine erythrocyte membrane skeleton. Sci Rep 11. doi:10.1038/s41598-021-88902-4

Prudêncio M, Rodrigues CD, Ataíde R, Mota MM. 2008. Dissecting in vitro host cell infection by Plasmodium sporozoites using flow cytometry. Cell Microbiol 10:218–24. doi:10.1111/j.1462-5822.2007.01032.x

Ramakrishnan C, Delves MJ, Lal K, Blagborough AM, Butcher G, Baker KW, Sinden RE. 2013. Laboratory maintenance of rodent malaria parasites. Methods Mol Biol 923:51–72. doi:10.1007/978-1-62703-026-7_5

Risco-Castillo V, Topçu S, Marinach C, Manzoni G, Bigorgne AE, Briquet S, Baudin X, Lebrun M, Dubremetz JF, Silvie O. 2015. Malaria sporozoites traverse host cells within transient vacuoles. Cell Host Microbe 18:593–603. doi:10.1016/j.chom.2015.10.006

Santos JM, Egarter S, Zuzarte-Luís V, Kumar H, Moreau CA, Kehrer J, Pinto A, da Costa M, Franke-Fayard B, Janse CJ, Frischknecht F, Mair GR. 2017. Malaria parasite LIMP protein regulates sporozoite gliding motility and infectivity in mosquito and mammalian hosts. Elife 6. doi:10.7554/eLife.24109

Sidik SM, Huet D, Ganesan SM, Huynh MH, Wang T, Nasamu AS, Thiru P, Saeij JPJ, Carruthers VB, Niles JC, Lourido S. 2016. A Genome-wide CRISPR Screen in Toxoplasma Identifies Essential Apicomplexan Genes. Cell 166:1423-1435.e12. doi:10.1016/j.cell.2016.08.019

Silvie O, Franetich JF, Boucheix C, Rubinstein E, Mazier D. 2007. Alternative invasion pathways for plasmodium berghei sporozoites. Int J Parasitol 37:173–182. doi:10.1016/j.ijpara.2006.10.005

Steinbuechel M, Matuschewski K. 2009. Role for the plasmodium sporozoite-specific transmembrane protein S6 in parasite motility and efficient malaria transmission. Cell Microbiol 11. doi:10.1111/j.1462-5822.2008.01252.x

Sultan AA, Thathy V, Frevert U, Robson KJH, Crisanti A, Nussenzweig V, Nussenzweig RS, Ménard R, Menard R, Ménard R. 1997. TRAP is necessary for gliding motility and infectivity of plasmodium sporozoites. Cell 90:511–522. doi:10.1016/S0092-8674(00)80511-5

Swearingen KE, Lindner SE, Flannery EL, Vaughan AM, Morrison RD, Patrapuvich R, Koepfli C, Muller I, Jex A, Moritz RL, Kappe SHI, Sattabongkot J, Mikolajczak SA. 2017. Proteogenomic analysis of the total and surface-exposed proteomes of Plasmodium vivax salivary gland sporozoites. PLoS Negl Trop Dis 11:e0005791. doi:10.1371/journal.pntd.0005791

Thompson J, Cooke RE, Moore S, Anderson LF, Janse CJ, Waters AP. 2004. PTRAMP; a conserved Plasmodium thrombospondin-related apical merozoite protein. Mol Biochem Parasitol 134. doi:10.1016/j.molbiopara.2003.12.003

Tibúrcio M, Yang ASP, Yahata K, Suárez-Cortés P, Belda H, Baumgarten S, van de Vegte-Bolmer M, van Gemert G-J, van Waardenburg Y, Levashina EA, Sauerwein RW, Treeck M. 2019. A Novel Tool for the Generation of Conditional Knockouts To Study Gene Function across the Plasmodium falciparum Life Cycle. MBio 10:e01170–19. doi:10.1128/mbio.01170-19

Yahata K, Hart MN, Davies H, Asada M, Wassmer SC, Templeton TJ, Treeck M, Moon RW, Kaneko O. 2021. Gliding motility of Plasmodium merozoites. Proc Natl Acad Sci U S A 118. doi:10.1073/pnas.2114442118

Yoshida N, Nussenzweig RS, Potocnjak P, Nussenzweig V, Aikawa M. 1980. Hybridoma produces protective antibodies directed against the sporozoite stage of malaria parasite. Science 207:71–73. doi:10.1126/science.6985745

Zhang M, Wang C, Otto TD, Oberstaller J, Liao X, Adapa SR, Udenze K, Bronner IF, Casandra D, Mayho M, Brown J, Li S, Swanson J, Rayner JC, Jiang RHY, Adams JH. 2018. Uncovering the essential genes of the human malaria parasite Plasmodium falciparum by saturation mutagenesis. Science 360. doi:10.1126/science.aap7847

